# Spatial collinearity constrains multivariate molecular-enriched network estimation

**DOI:** 10.64898/2026.06.10.731385

**Authors:** Timothy Lawn, Johan Nakuci, Steve CR Williams, Federico Turkheimer, Mitul A. Mehta

**Affiliations:** Oxford Centre for Integrative Neuroimaging, Department of Experimental Psychology, University of Oxford; Department of Neuroimaging, Institute of Psychiatry, Psychology and Neuroscience, King’s College London; Athinoula A. Martinos Center for Biomedical Imaging, Massachusetts General Hospital and Harvard Medical School; U.S. Army DEVCOM Army Research Laboratory

**Keywords:** REACT, Networks, PET, fMRI, Molecular, Collinearity, Pharmacoimaging

## Abstract

Analyses of neuroimaging data increasingly leverage the distribution of neurotransmitter receptors derived from Positron Emission Tomography (PET) to bridge the gap between micro- and macro-scale brain function. However, these receptor maps are highly spatially overlapping which can give rise to interpretive and analytical challenges. Here, we systematically investigate the impact of spatial collinearity among PET maps in the context of Receptor-Enriched Analysis of functional Connectivity by Targets (REACT), a method that uses receptor maps as spatial regressors to derive subject-level molecular-enriched functional connectivity networks. Exhaustive combinatorial analysis across 19 receptor and transporter maps showed that collinearity scales rapidly with the number of receptors modelled simultaneously, and that this was relatively stable across parcellation scales, reflecting the intrinsic organisation of neurotransmitter systems. Using test-retest fMRI data from the Human Connectome Project, we demonstrate that modelling greater numbers of receptors degrades the reliability of molecular-enriched networks derived from conventional multivariate REACT models, and that collinearity among receptor maps drives this degradation. An alternative univariate approach, in which each receptor is modelled independently, yielded more reliable networks and, when applied to a within-subjects study of LSD compared to placebo, better recovered the role of the 5HT-2A receptor in LSD’s neural effects. These findings identify spatial collinearity as a fundamental constraint on multivariate molecular-enriched network estimation and support univariate modelling as a more robust default for this class of analysis.

## Introduction

The blood-oxygen level dependent (BOLD) signal measured by functional magnetic resonance imaging (fMRI) provides no intrinsic specificity for the molecular mechanisms that give rise to it (Hillman, 2014; Lawn et al., 2023a). This disconnect has left pharmaco-fMRI studies conceptually detached from the molecular targets through which drugs act. Recent advances in multimodal imaging have sought to bridge this gap by leveraging PET-derived spatial distributions of neurotransmitter receptors and transporters to inform fMRI analyses (Bazinet et al., 2023; Dukart et al., 2021; Hansen and Misic, 2025; Lawn et al., 2023a; Markello et al., 2022; Royer et al., 2026). However, because neurotransmitter systems are not spatially independent, these approaches are vulnerable to the inferential and analytical challenges associated with high levels of spatial similarity (collinearity) among receptor maps.

Among these approaches, Receptor-Enriched Analysis of functional Connectivity by Targets (REACT; (Dipasquale et al., 2019)) has emerged as a widely used method for disentangling pharmacological mechanisms (Boucherie et al., 2023; Dipasquale et al., 2020; Lawn et al., 2023b, 2022; Shatalina et al., 2025; van den Bosch and Cools, 2026) and predicting treatment response (Martins et al., 2022; Tolle et al., 2025). REACT uses PET-derived receptor density maps as spatial templates in a dual-regression framework (Nickerson et al., 2017) to extract subject-specific functional networks tied to each neurotransmitter system (Dipasquale et al., 2019). Conventionally, all receptors of interest are entered simultaneously into both spatial and temporal regression stages, with the intention of isolating variance unique to each receptor by partialling out the contributions of all others (Dipasquale et al., 2019) (see (Lawn et al., 2023a) for review). This multivariate strategy assumes that the receptor maps are sufficiently spatially distinct to permit stable estimation of their individual contributions, yet unlike the spatially independent networks (Beckmann and Smith, 2004) for which dual regression was originally designed (Nickerson et al., 2017), neurotransmitter receptor maps are not orthogonal.

Multiple receptors and transporters show correlated distributions across the brain, reflecting neurotransmitter co-release (Vaaga et al., 2014), interactions (Avery and Krichmar, 2017), and overlapping functional roles (Froudist-Walsh et al., 2023; Hansen et al., 2022). When two receptor PET maps share substantial spatial variance, REACT’s regression models may not stably attribute the shared portion to either one, inflating standard errors and making coefficient estimates sensitive to minor perturbations in the data or model specification. Beyond estimation instability, spatial collinearity creates a broader inferential problem: when receptor maps are highly correlated, a strong spatial correspondence between a brain map and a given receptor distribution cannot be unambiguously attributed to that receptor, a confound that extends to the full class of brain map contextualisation analyses (Royer et al., 2026). These concerns are not merely theoretical. (van den Bosch and Cools, 2026) recently found that substantial spatial overlap between dopamine and serotonin transporter maps precluded disentangling their known roles in the pharmacology of methylphenidate using REACT (see (Lawn and Mehta, 2026) for commentary). Furthermore, they observed attenuated drug effects when receptors were modelled together in a multivariate REACT analysis compared with independent univariate models for each receptor (van den Bosch and Cools, 2026). Despite this, the collinearity landscape across available receptor maps and its consequences for REACT remain largely unexplored.

Here, we systematically investigate how spatial collinearity affects molecular-enriched network estimation. First, we characterise the collinearity landscape across 19 receptor and transporter maps, examining how it scales with model complexity and parcellation resolution. Second, using test-retest fMRI data, we examine whether collinearity degrades the reliability of multivariate molecular-enriched network estimation and test whether a univariate alternative - in which each receptor is modelled independently rather than simultaneously - yields more reliable networks. The univariate approach eliminates inter-receptor competition for shared spatial variance at the cost of forgoing subject-level adjustment for other systems. We test this trade-off empirically. Third, we apply both modelling approaches to an example pharmacoimaging dataset where we selected LSD to test whether the univariate pipeline better recovers the established role for 5HT-2A receptors in psychedelics action. Together, these analyses demonstrate how collinearity constrains multivariate molecular-enriched network estimation and test whether univariate modelling offers a more robust alternative.

## Methods

### PET Receptor and Transporter Maps

Volumetric PET receptor and transporter density maps were obtained from the Hansen atlas ((Hansen et al., 2022); https://github.com/netneurolab/hansen_receptors) which aggregates normative PET data from over 1,200 healthy individuals across multiple radiotracer studies. Maps for 19 neurotransmitter targets were included, spanning the serotonergic (5HT-1A, 5HT-1B, 5HT-2A, 5HT-4, 5HT-6, 5HTT), dopaminergic (D1, D2, DAT), glutamatergic (mGluR5, NMDA), GABAergic (GABA-A), noradrenergic (NAT), cholinergic (α4β2, M1, VAChT), histaminergic (H3), cannabinoidergic (CB1), and opioidergic (MU) systems.

All maps are provided in the MNI152 standard template space but at varying native resolutions. Those not already at 2mm^3^ isotropic resolution (91×109×91 voxels) were resampled using ANTs (antsApplyTransforms) with linear interpolation (Avants et al., 2011). Maps were parcellated using combined cortical and subcortical atlases at ten spatial scales. Cortical parcels were defined by the Schaefer 7-network parcellation (Schaefer et al., 2018) at scales of 100 to 1,000 parcels (in increments of 100), each combined with a corresponding scale of the Tian subcortical atlas (Tian et al., 2020): S1 (16 parcels) for the 100- and 200-parcel cortical schemes, S2 (32 parcels) for the 300-to 500-parcel schemes, S3 (50 parcels) for the 600- and 700-parcel schemes, and S4 (54 parcels) for the 800-to 1,000-parcel schemes, yielding total parcel counts ranging from 116 to 1,054. For each parcellation, mean PET values were computed within each parcel after excluding zero-valued voxels. This multi-scale approach enabled examination of how spatial resolution affects collinearity. When not explicitly stated otherwise, subsequent analyses used the Schaefer 400 + Tian S2 scheme (432 parcels) as the standard reference parcellation.

Twelve of the 19 receptor/transporter targets were represented by multiple independent tracer studies in the Hansen atlas (ranging from 2 to 4 maps per target). For these, consensus maps were derived following the procedure described in (Hansen et al., 2022): each individual parcellated map was first z-scored across parcels to place all tracer studies on a common scale, then a weighted average was computed using the number of healthy control participants contributing to each study as weights. The resulting consensus map was min-max scaled to the 0-1 range. For targets represented by a single tracer study, parcellated values were min-max scaled to the 0-1 range directly.

### Characterisation of Spatial Collinearity

Pairwise Pearson correlations were computed across all 19 maps and ordered by hierarchical clustering (Ward’s method) to expose the block structure of inter-receptor spatial similarity.

We next performed an exhaustive combinatorial analysis to characterise collinearity among these PET maps. Variance inflation factors (VIF) were computed to quantify collinearity for every possible combination of k receptor maps (k=1-19), yielding C(19, k) combinations at each model size. VIFs range from 1 to ∞, with lower values reflecting lower levels of collinearity (O’Brien, 2007). The VIF for the *j*^th^ receptor in a given model is defined as:

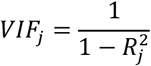

where 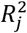 is the coefficient of determination obtained by regressing receptor map *j* on all other receptor maps in the model. For each combination, we summarised the collinearity landscape by examining: (i) the distribution of mean VIF (averaged across all receptors within each combination) at each model size, (ii) the percentage of combinations in which at least one receptor exceeded common thresholds for moderate (VIF>5) or severe (VIF>10) collinearity, using the maximum individual VIF within each combination (VIF_max_), and (iii) per-receptor mean VIF as a function of k, to identify which receptor maps are most susceptible to collinearity. Thresholds of 5 and 10 are common heuristics for moderate and severe multicollinearity (García et al., 2015; Kim, 2019; Kovàcs et al., 2005) and are used here as descriptive benchmarks rather than formal inferential cutoffs (O’Brien, 2007) as in several recent REACT papers (Lawn et al., 2024, 2023b; Lawn and Mehta, 2026). Principal component analysis (PCA) was also applied to the z-scored 19-receptor design matrix to characterise its effective dimensionality.

To assess the influence of spatial scale, the full combinatorial VIF analysis and pairwise correlation matrix were computed at each of the ten parcellation scales (116-1,054 parcels). The stability of the inter-receptor correlation structure across scales was quantified by computing the Pearson correlation between the vectorised upper-triangular elements of each pair of scale-specific correlation matrices. Mean VIF was examined as a joint function of model size (k) and parcellation scale to determine whether particular resolutions offer more or less favourable conditions for multivariate modelling.

### REACT Pipelines

Molecular-enriched functional connectivity networks were estimated using a parcellated implementation of REACT. REACT employs a dual-regression framework (Nickerson et al., 2017) in which PET-derived receptor density maps serve as spatial priors to extract subject-specific functional networks associated with each neurotransmitter system (Dipasquale et al., 2019). In the first stage (spatial regression), BOLD values across parcels at each timepoint are regressed onto the PET receptor maps, yielding one coefficient per receptor per timepoint. Concatenating these across timepoints produces a receptor-weighted timeseries for each map in the model. In the second stage (temporal regression), these receptor-weighted timeseries are regressed against the fMRI data at each parcel across timepoints, yielding a spatial map of coefficients for each receptor that represents the subject-specific molecular-enriched functional connectivity network. Prior to regression, PET maps were demeaned across parcels and fMRI data were demeaned at each timepoint (stage 1) or across timepoints at each parcel (stage 2). In stage 2, the receptor-weighted timeseries were additionally standardised to unit variance. Parcels with all-zero fMRI signal were excluded from both stages and assigned zero values in the output maps. For more information on REACT, see review (Lawn et al., 2023a).

In the multivariate REACT pipeline, all receptor maps were entered simultaneously as columns in the first-stage spatial regression, and the resulting set of receptor-weighted timeseries were entered simultaneously as temporal regressors in the second stage. Both regression stages are therefore multivariate: the first-stage timeseries for each receptor reflect BOLD fluctuations associated with that receptor’s spatial distribution after partialling out all other receptor maps, and the second-stage spatial maps further partial out temporal variance shared among the receptor-weighted timeseries. In the univariate pipeline, each receptor map was modelled independently through its own first-stage spatial regression and its own second-stage temporal regression, with no other receptor entering either stage. This eliminates inter-receptor competition for shared variance at both stages, such that each receptor’s FC map captures all co-distributed BOLD variance regardless of overlap with other systems. Because the regression coefficients themselves constitute the core output of REACT (each coefficient map represents a subject-specific molecular-enriched network) the relevant criterion for evaluating pipeline performance is the stability and interpretability of these coefficients, not the overall variance explained by the model. This distinguishes REACT from predictive regression contexts in which model fit is the primary objective and coefficient instability is tolerable.

### HCP Test-Retest Dataset

Resting-state fMRI data were obtained from the Human Connectome Project Young Adult (HCP-YA) test-retest dataset (Glasser et al., 2013; Van Essen et al., 2012). This dataset comprises 45 participants who underwent the full HCP 3T imaging protocol on two separate occasions. All participants provided written informed consent, and the study was approved by the Washington University Institutional Review Board (IRB number 201204036).

Data were acquired on a customised Siemens 3T Connectome Skyra scanner with a 32-channel head coil using a gradient echo EPI sequence with multiband acceleration (TR=720ms, TE=33.1ms, flip angle=52°, 2mm^3^ voxels, 72 slices, multiband factor=8, 1,200 frames per run, eyes open with fixation). At each visit, four resting-state runs (two phase-encoding directions) were acquired in each of two sessions. We used fully processed concatenated session data (rfMRI_REST_Atlas_MSMAll_hp2000_clean_rclean_tclean.dtseries.nii) from the HCP-YA 2025 release, which includes updated preprocessing relative to the S1200 release (https://balsa.wustl.edu/project?project=HCP_Retest for details). BOLD timeseries were extracted by averaging vertices (cortical) or voxels (subcortical) within each parcel of the Schaefer 400 + Tian S2 atlas in CIFTI grayordinates space, yielding a 432-parcel × timepoint matrix per run.

For each participant, molecular-enriched FC maps were estimated using both univariate and multivariate REACT pipelines described above. Test-retest reliability was quantified at the parcel level using the intraclass correlation coefficient (ICC(3,1)) between sessions across participants. ICC(3,1) was selected because scanning sessions are fixed levels (the same two visits are compared across all participants)(Koo and Li, 2016). To systematically examine how model complexity affects reliability, the multivariate pipeline was applied to every possible combination of k receptor maps (k=2-19), yielding C(19, k) models at each model size, and the univariate pipeline provided the k=1 baseline for each receptor individually. Mean ICC was computed for each receptor within each combination and compared across model sizes. The significance of the reliability decline was assessed using a within-receptor permutation test (10,000 permutations). Mean ICC values were aggregated across all combinations at each k for each receptor, yielding a receptor × k matrix. The test statistic was the mean within-receptor slope of ICC on k; the null distribution was generated by independently permuting ICC values across k-levels within each receptor.

To test whether collinearity itself - rather than model size per se - predicted the magnitude of reliability degradation, per-receptor ICC and VIF values were first averaged across all combinations at each k-level, yielding one observation per receptor per k (19 receptors × 18 k-levels = 342 cells). This aggregation eliminates pseudo-replication from overlapping receptor combinations at each model size. A linear mixed-effects model was then fitted (using statsmodels in Python) with receptor as a random intercept: ΔICC ∼ log_10_(VIF) + (1 | receptor), where ΔICC is the change in ICC relative to the univariate baseline. A sensitivity model additionally included k as a fixed effect to assess whether VIF remained predictive after adjusting for model size. Nested models were compared using likelihood ratio tests and Bayesian information criteria (BIC).

### LSD Pharmacoimaging Dataset

We performed secondary analyses of an openly available (https://openneuro.org/datasets/ds003059/versions/1.0.0) pharmacoimaging dataset (Carhart-Harris et al., 2016). The original study was approved by the National Research Ethics Service committee London-West London and conducted in accordance with the Declaration of Helsinki (2000). Fifteen healthy volunteers completed a within-subjects, placebo-controlled, balanced-order design. Participants received either LSD (75µg intravenous) or placebo (saline), with sessions separated by at least two weeks. Resting-state fMRI data (eyes closed) were acquired approximately 70 minutes post-dosing on a 3T GE HDx system (TR/TE=2,000/35ms, 3.4mm isotropic, 7-minute runs). Three runs per session were acquired; runs 1 and 3 (resting-state) were analysed here, excluding run 2 (music listening). fMRI data were preprocessed as described in (Carhart-Harris et al., 2016) and parcellated using the Schaefer 400 + Tian S2 scheme (432 parcels).

Five receptor maps were selected based on LSD’s known receptor pharmacology: 5HT-1A, 5HT-2A, 5HTT, D1, and D2. LSD’s subjective and neural effects are largely abolished by 5HT-2A antagonism (Preller et al., 2018), establishing 5HT-2A as the primary pharmacological driver. At the circuit level, this is thought to operate through 5HT-2A-mediated modulation of pyramidal cell gain (Burt et al., 2021), which in turn promotes the large-scale network state transitions observed under LSD (Singleton et al., 2022). LSD is therefore a useful test case where a valid method would be expected to recover 5HT-2A-dominant drug effects. Furthermore, the availability of multiple runs per session offered the crucial opportunity to examine reliability. Molecular-enriched FC maps were estimated for each participant, session (LSD and placebo), and run (1 and 3) using both the univariate and multivariate pipelines with all five receptor maps.

Within-session reliability was quantified as the Spearman correlation between FC maps from runs 1 and 3. This was computed separately within each session (LSD and placebo) and then averaged across sessions for each participant, receptor, and pipeline, yielding a single session-general reliability estimate per subject. Between-pipeline differences were tested via sign-flipping permutation tests (10,000 permutations). Permutation p-values were corrected for multiple comparisons across five receptors using the Benjamini-Hochberg false discovery rate procedure. Spearman correlation was preferred over ICC given the limited sample size (N=15). Group-level drug effects were assessed using paired permutation tests (LSD versus placebo) at each parcel for each receptor network and pipeline using networks averaged across runs (1 and 3). Parcel-wise permutation p-values (10,000 sign-flipping permutations) were corrected using the Benjamini-Hochberg false discovery rate (FDR; q<0.05) (Benjamini and Hochberg, 1995), with a further Bonferroni correction across the five receptor maps (i.e., FDR applied at q < 0.05/5 = 0.01 per receptor).

To assess receptor specificity, the absolute parcel-wise t-statistics from the 5HT-2A network were compared against those from each comparator receptor network. For each pairwise comparison, we computed the summed absolute t-statistic mass favouring 5HT-2A versus the comparator (i.e. the sum of |t| differences across parcels where 5HT-2A showed the larger effect, and vice versa), expressed as a proportion. Additionally, each parcel was assigned to the receptor with the largest absolute drug-effect t-statistic (receptor dominance), and the redistribution of dominant receptor assignments between the univariate and multivariate pipelines was characterised using Sankey diagrams.

## Results

### Spatial collinearity among PET receptor maps scales with model complexity

The 19 PET receptor and transporter maps showed substantial spatial collinearity, with hierarchical clustering exposing blocks of highly correlated receptors within and across neurotransmitter systems (Figure 2A). Pairwise Pearson correlations ranged from r=-0.59 to r=0.95 (median r=0.28), with 31% of the 171 unique pairs exceeding

**Figure 1.**
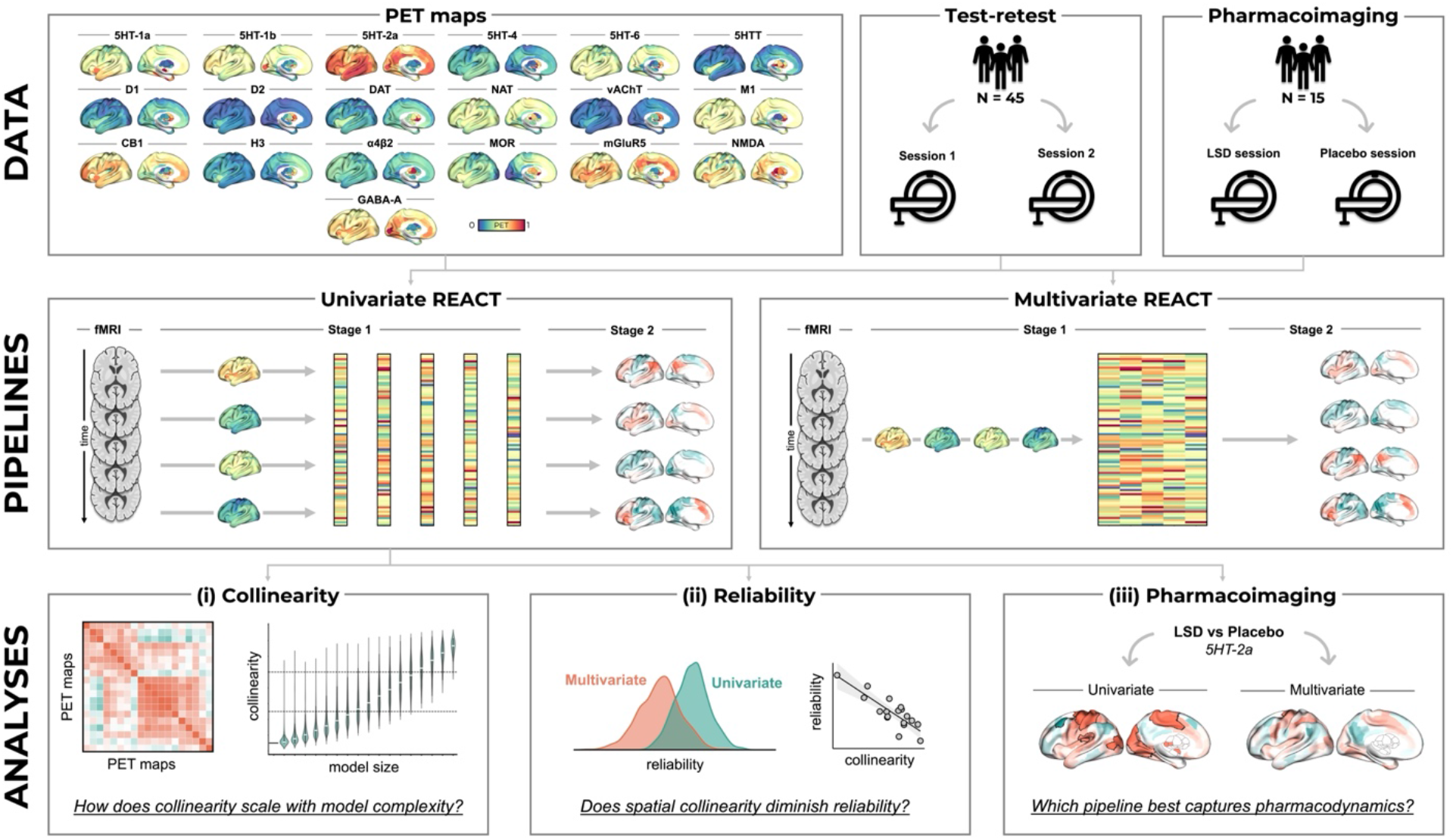
Study overview. Three rows outline the data, pipelines, and analyses. **Top (Data)**: Nineteen PET receptor and transporter maps from the Hansen atlas, parcellated using combined Schaefer cortical and Tian subcortical atlases (left hemisphere maps shown at the Schaefer 400 + Tian S2 scale; 432 parcels). Two fMRI datasets were analysed: resting-state test-retest data from the Human Connectome Project (N = 45, two visits) and a within-subjects LSD pharmacoimaging dataset (N = 15, LSD and placebo sessions). **Middle (Pipelines)**: In the univariate pipeline (left), each receptor map is modelled independently through its own spatial regression (Stage 1: BOLD timeseries regressed onto a single PET map) and temporal regression (Stage 2: the resulting receptor-weighted timeseries regressed against BOLD data at each parcel), yielding one molecular-enriched FC network per receptor. In the multivariate pipeline (right), all receptor maps enter Stage 1 simultaneously as columns of a single design matrix, and the resulting set of receptor-weighted timeseries enter Stage 2 simultaneously, such that both stages partial out variance shared among receptors. **Bottom (Analyses)**: Three empirical components: **(i)** exhaustive combinatorial VIF analysis characterising how spatial collinearity among PET maps scales with model size; **(ii)** test-retest reliability analysis examining whether collinearity degrades the reliability of multivariate molecular-enriched networks; **(iii)** LSD pharmacoimaging analysis testing whether the univariate pipeline better recovers the established 5HT-2A receptor specificity of psychedelics.

**Figure 2.**
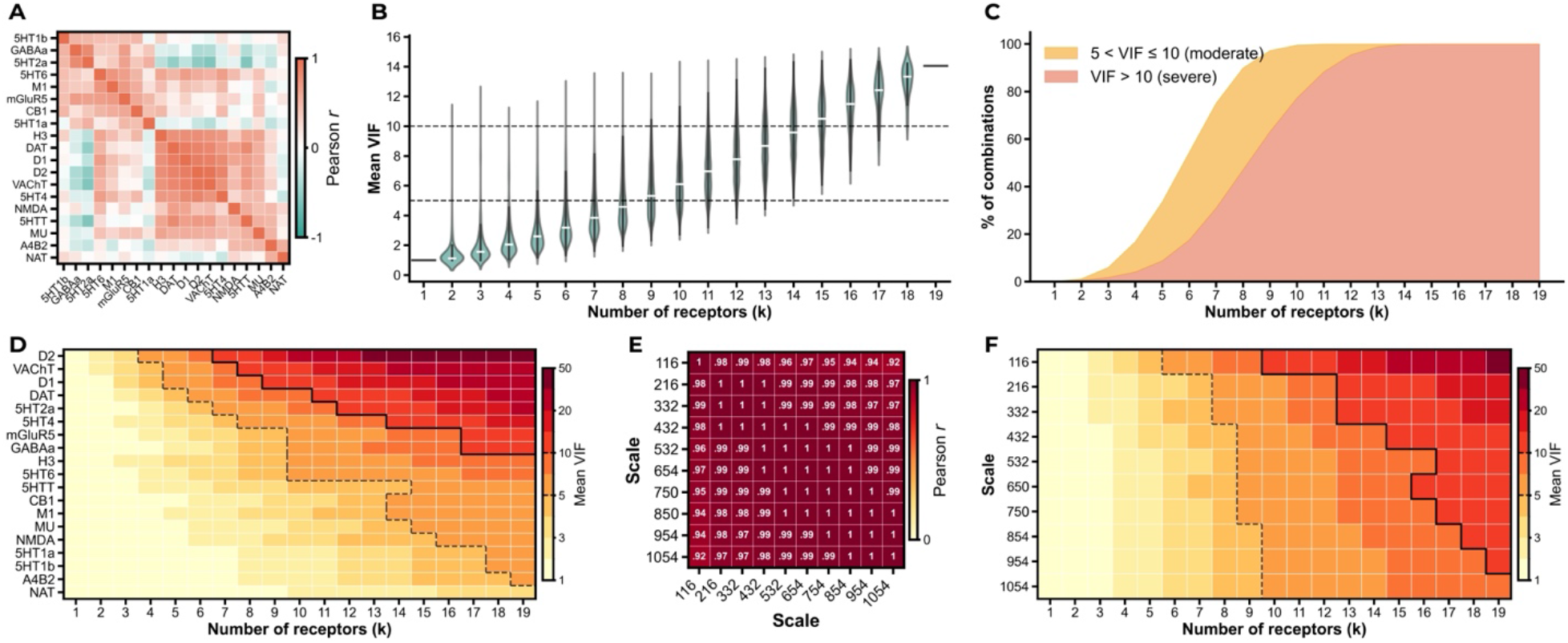
Spatial collinearity across PET map combinations and parcellation scales. All panels use the Schaefer 400 + Tian S2 parcellation (432 parcels) unless otherwise stated. **(A)** Pearson correlation matrix of the 19 PET receptor and transporter maps hierarchically clustered using Ward’s method. **(B)** Distribution of mean VIF values (averaged across all receptors within each combination) across all C(19, k) receptor combinations at each model size (k=1–19). Dashed and solid staircase boundaries trace VIF=5 and VIF=10, respectively. **(C)** Percentage of receptor combinations in which at least one receptor exceeds moderate (5<VIF ≤ 10; orange) or severe (VIF>10; red) collinearity thresholds (VIF_max_), as a function of model size. **(D)** Mean VIF per receptor across all combinations at each model size, ordered by mean VIF across all k (highest at top). Dashed and solid staircase boundaries trace where mean VIF exceeds 5 and 10, respectively. **(E)** Stability of the inter-receptor correlation structure across ten parcellation scales (116–1,054 parcels). Each cell shows the Pearson correlation between the vectorised upper triangles of the scale-specific inter-receptor correlation matrices. **(F)** Mean VIF across all receptor combinations as a joint function of model size (k) and parcellation scale. Dashed and solid staircase boundaries trace VIF=5 and VIF=10, respectively.

|r|>0.5. PCA confirmed that collinearity compresses the effective dimensionality of receptor space: four components accounted for 80% of the variance and nine for 95% (Supplementary Figure 1), indicating far fewer independent dimensions than the 19 targets suggest.

Exhaustive combinatorial VIF analysis across all C(19, k) receptor combinations demonstrated that collinearity scales rapidly with model size (Figure 2B). The median mean VIF first exceeded the moderate threshold of 5 at k=9 receptors and the severe threshold of 10 at k=15. Substantial variability existed across combinations at each model size, with at least one combination exceeding a mean VIF of 10 at every k≥2. When considering VIF_max_ (the highest individual VIF within a combination), the majority of combinations exceeded moderate collinearity (>5) by k=6 and severe collinearity (>10) by k=9 (Figure 2C).

In the full 19-receptor model, every receptor except NAT (VIF=3.6) exceeded VIF=5, eight exceeded VIF=10, and the worst-affected receptor (D2) reached VIF=59.7. Per-receptor VIF profiles revealed a clear hierarchy of collinearity susceptibility (Figure 2D). The most affected receptors, D2, VAChT, and D1 (mean VIF across all k: 21.1, 13.6, and 12.7), are those whose spatial distributions are most strongly predicted by other maps in the set. Conversely, NAT, 5HT-1A, and α4β2 demonstrated more spatially distinctive distributions.

The inter-receptor correlation structure was remarkably stable across parcellation scales (Figure 2E). All pairwise correlations between scale-specific correlation matrices exceeded r=0.92 (mean r=0.98), indicating that the collinearity architecture reflects the intrinsic organisation of neurotransmitter systems rather than resolution-dependent artefact. Mean VIF as a joint function of model size and parcellation scale (Figure 2F) confirmed that the number of receptors was the dominant driver of collinearity, though the coarsest parcellations (116–216 parcels) modestly amplified it at large k, consistent with spatial averaging increasing inter-map correlations.

### Multivariate modelling degrades the reliability of molecular-enriched networks

We next tested whether this collinearity has measurable consequences for the reliability of REACT-derived networks. Reliability declined monotonically with the number of receptors in the multivariate model (Figure 3A). Mean ICC fell from 0.547 (k=1, univariate) to 0.448 (k=19, full model), an 18% reduction that, by conventional benchmarks (Cicchetti, 1994), shifts the full model from the upper to the lower boundary of fair reliability (0.40–0.59), approaching the threshold for poor reliability.

**Figure 3.**
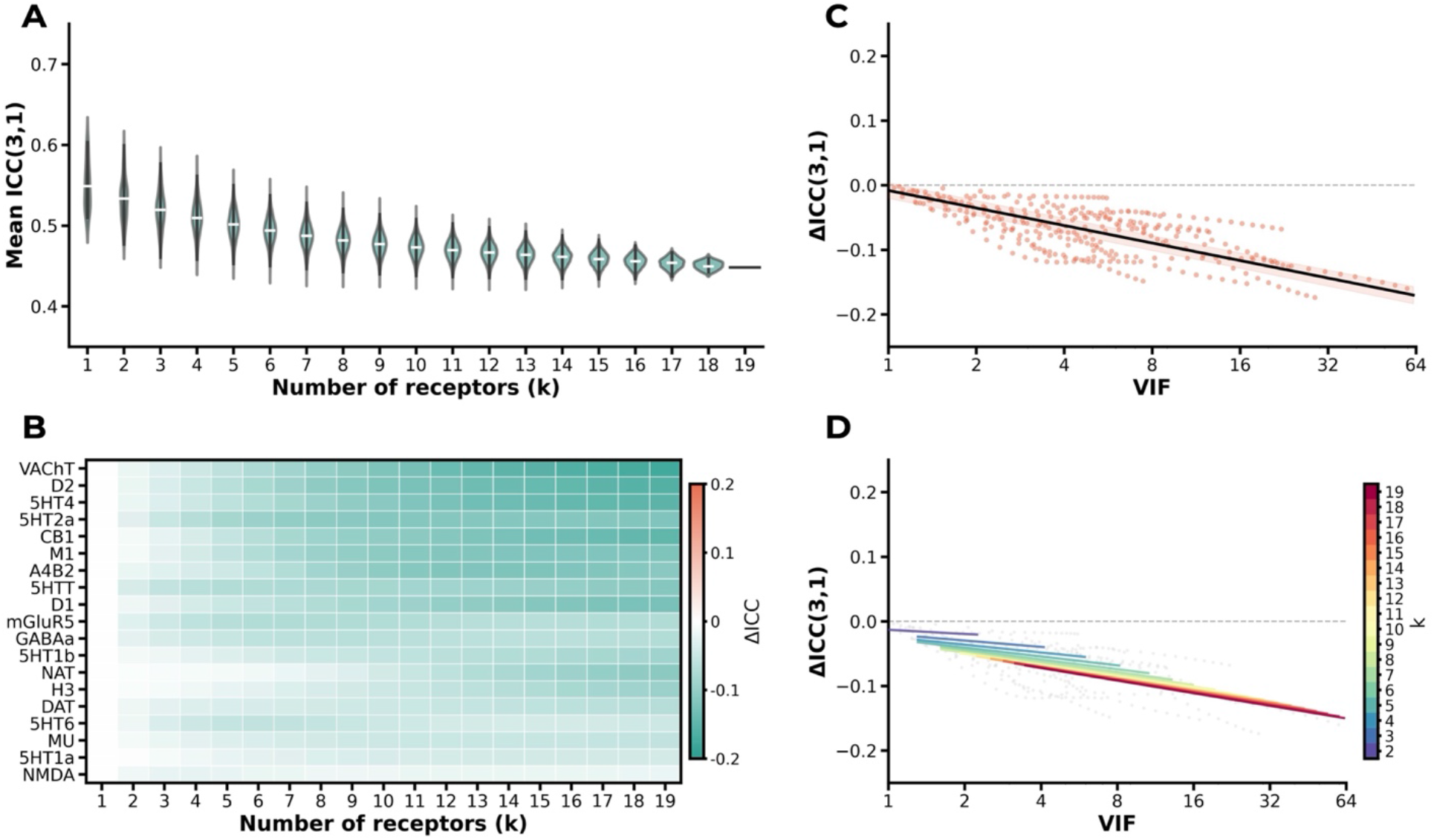
Multivariate modelling degrades REACT network reliability. **(A)** Distribution of mean ICC(3,1) values across all receptor combinations at each model size (k). At k=1, each point is one receptor’s mean ICC from the univariate pipeline (n=19 receptors); at k=2– 18, violins span all C(19, k) multivariate combinations; at k=19, the single full model is shown. White bars indicate the median. **(B)** ΔICC (multivariate minus univariate baseline) per receptor as a function of model size (k). Receptors are ordered by mean ΔICC across k-levels (most affected at top). **(C)** Collinearity predicts reliability loss. Each point represents one receptor at one k-level (19 receptors × 18 k-levels = 342 cells), with both ΔICC and VIF averaged across all combinations at that k-level. The fitted line and shaded 95% confidence interval are from a linear mixed-effects model with receptor as a random intercept. **(D)** The VIF-ΔICC relationship holds within each model size. Each line is a univariate OLS fit at a single k-level (k=2-19), with colour indicating model size. Grey points show the 342 receptor × k-level observations.

A within-receptor permutation test confirmed this decline (mean slope=−0.005, p=0.0001, 10,000 permutations). The decline was not uniform across receptors, with receptors such as D2 and VAChT, whose spatial distributions overlap extensively with other maps, showing the steepest drops (Figure 3B). Conversely, the NMDA and 5HT-1A receptors occupy more spatially distinctive distributions and were relatively protected. This heterogeneity suggested that the degree of collinearity a given receptor experiences - rather than model size alone - determines the magnitude of reliability loss. The collinearity susceptibility hierarchy identified in Figure 2D further predicted the magnitude of reliability degradation at the receptor level (Spearman ρ=−0.50, p=0.028, n=19; Supplementary Figure 2).

Per-receptor ICC and VIF values were aggregated across all combinations at each k-level (19 receptors × 18 k-levels = 342 cells), eliminating pseudo-replication, and linear mixed-effects models were fitted with receptor as a random intercept. Higher collinearity strongly predicted reliability loss (b=−0.090, SE=0.003, p<0.0001; Figure 3C). To disentangle VIF from model size, three nested models were compared via likelihood ratio tests on maximum-likelihood fits. Adding k to the VIF model did not improve fit (LRT χ^2^=1.83, p=0.176, BIC: VIF-only −1783 vs VIF+k −1779), and k was not significant when VIF was already included (b=−0.001, SE=0.001, p=0.184). Conversely, adding VIF to a k-only model yielded a substantial improvement (LRT χ^2^=66.15, p<0.0001). This asymmetry indicates that collinearity carries information about reliability degradation beyond what model size explains, while model size adds little beyond collinearity. Within-k Spearman correlations between VIF and ΔICC ranged from ρ=−0.36 to ρ=−0.60 (Figure 3D), with the strongest associations at intermediate k where between-receptor variance in collinearity is greatest.

### Collinearity compromises pharmacological specificity in LSD pharmacoimaging

We next examined whether these effects extend to a typical pharmacoimaging application of REACT. The two pipelines yielded networks with overlapping but distinct spatial topographies (Figure 4A), with subject-level Spearman correlations between their FC maps ranging from ρ=0.48 to ρ=0.90 across receptor–subject combinations (Figure 4B). Even within this focused five-receptor model, VIF varied substantially across receptors (Figure 4C). The dopaminergic receptors D2 (VIF=7.2) and D1 (VIF=6.4) crossed the conventional moderate threshold, reflecting their overlapping striatal distributions, while the three serotonergic targets remained below it (5HT-1A=1.5, 5HTT=2.5, 5HT-2A=2.5). This was a relatively favourable design relative to the broader VIF landscape characterised in Figure 2.

**Figure 4.**
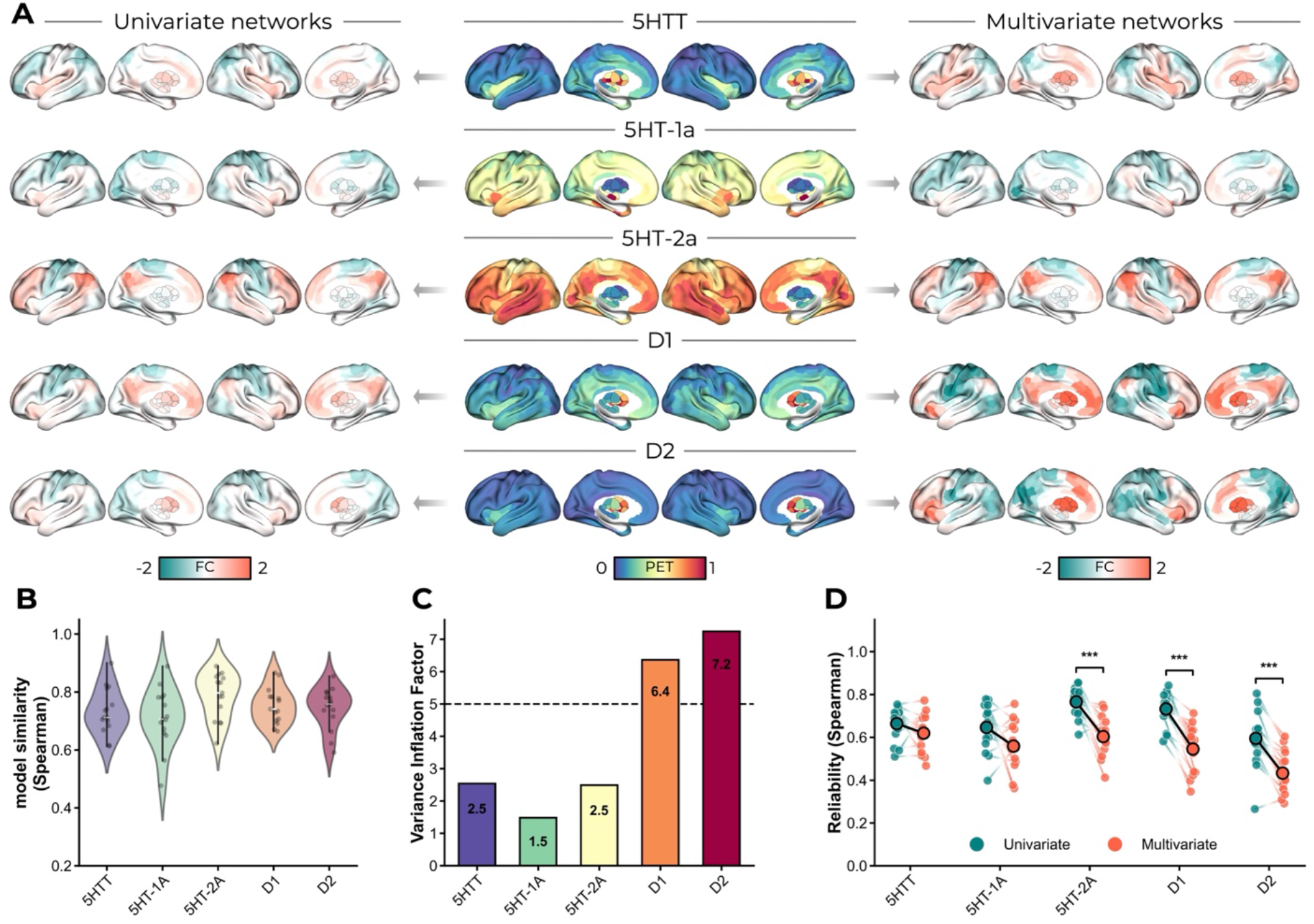
Collinearity and reliability in the LSD five-receptor model. N=15 participants and Schaefer-Tian 432 parcellation. **(A)** Group-mean molecular-enriched functional connectivity networks derived from the univariate (left) and multivariate (right) pipelines for each of the five receptor maps (centre). Insets show subcortical structures in medial view. **(B)** Spatial similarity between univariate and multivariate FC maps for each receptor, quantified as subject-level Spearman correlations. Each dot represents one participant and violins show the distribution across participants. **(C)** Variance inflation factors for the five-receptor multivariate model. Dashed line indicates the moderate threshold (VIF=5). **(D)** Within-session test–retest reliability (Spearman correlation between FC maps from independent runs) for the univariate (teal) and multivariate (red) pipelines. Individual participants are shown as paired dots connected by lines; large dots with black borders indicate group means. Significance brackets denote permutation tests (*** p<0.001, ** p<0.01, * p<0.05).

Despite this comparatively modest collinearity, within-session test-retest reliability was consistently higher for the univariate pipeline (Figure 4D). The univariate model yielded mean reliability values ranging from ρ=0.596 (D2) to ρ=0.766 (5HT-2A), compared with ρ=0.433 (D2) to ρ=0.620 (5HTT) for the multivariate model. Permutation tests confirmed significantly higher within-session reliability for the univariate pipeline for 5HT-2A (Δρ=0.162, p=0.0004, p_FDR_=0.0007), D1 (Δρ=0.186, p=0.0001, p_FDR_=0.0005), and D2 (Δρ=0.163, p=0.0003, p_FDR_=0.0007), with a trend for 5HT-1A (Δρ=0.088, p=0.065, p_FDR_=0.082). 5HTT did not show any difference between pipelines (Δρ=0.044, p=0.155, p_FDR_=0.155).

5HT-1A was the receptor whose spatial distribution overlaps least with the others in this set and showed no significant difference. The two receptors with the largest reliability gaps (D1, D2) were also the only two crossing the moderate VIF threshold, consistent with collinearity contributing to reliability degradation. However, 5HT-2A showed a comparable Δρ despite a much lower within-model VIF. This suggests that VIF for any specific receptor captures only part of what determines reliability under multivariate REACT, and that even modest model collinearity is sufficient to produce meaningful estimation instability.

### Multivariate modelling obscures 5HT-2A specificity of LSD effects

Parcel-wise paired t-tests (LSD versus placebo) revealed a striking dissociation between pipelines (Figure 5A). The univariate model identified 115 of 432 parcels (26.6%) with significant drug effects in the 5HT-2A network only. The multivariate model detected no significant effects in any receptor network (0 of 2,160 parcel-receptor tests survived correction). This complete null is consistent with the inflated variance and reduced reliability documented above: multivariate partialling attenuates effect sizes to the point where genuine pharmacological signals fall below the detection threshold.

**Figure 5.**
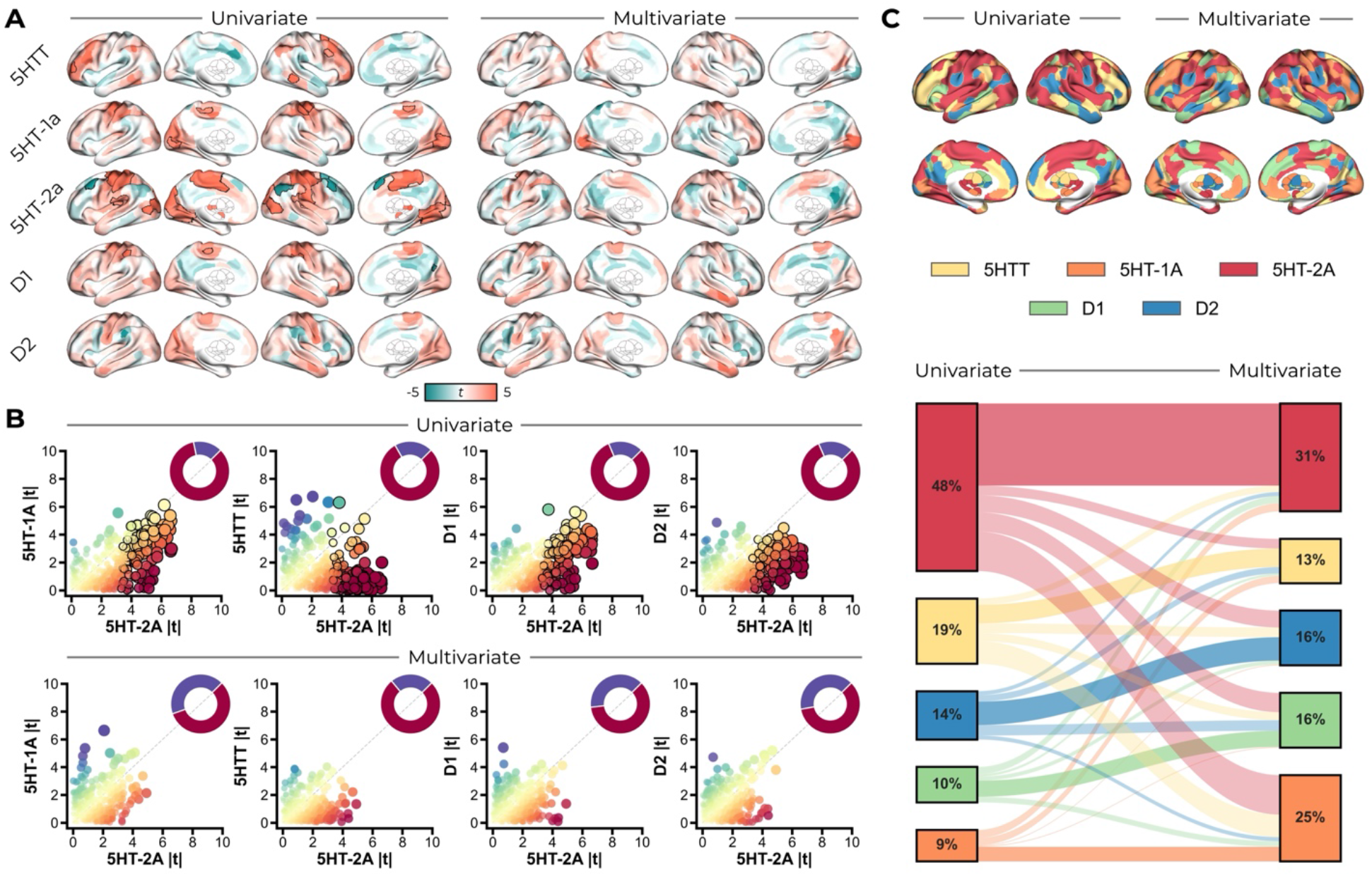
The univariate pipeline recovers 5HT-2A-dominant LSD effects that the multivariate pipeline fails to detect. **(A)** Parcel-wise paired t-statistics (LSD minus placebo) for each receptor network under the univariate (left) and multivariate (right) pipelines. Black outlines denote parcels surviving FDR correction (q < 0.01 per receptor, Bonferroni-corrected across five receptors). **(B)** Receptor specificity of LSD effects. Each column compares absolute parcel-wise t-statistics for 5HT-2A (x-axis) against a comparator receptor (y-axis) under the univariate (top row) and multivariate (bottom row) pipelines. Parcels below the diagonal show stronger effects for 5HT-2A. Point colour encodes the |t| difference (red: 5HT-2A dominant; blue: comparator dominant) and point size scales with the larger |t|. Black-outlined points survive FDR correction for at least one receptor in the pair. Inset donut charts show the proportion of summed |t| mass favouring 5HT-2A (red) versus the comparator (blue). **(C)** Receptor dominance analysis. Top: each parcel is assigned to the receptor with the largest absolute drug-effect t-statistic and coloured accordingly for the univariate (left) and multivariate (right) pipelines. Bottom: Sankey diagram tracing parcel-level transitions in dominant receptor assignment between pipelines; node height and percentages indicate the proportion of parcels dominated by each receptor and flow width is proportional to the number of parcels transitioning between dominant receptors.

To quantify 5HT-2A specificity across all parcels, we compared absolute parcel-wise t-statistics for the 5HT-2A network against each comparator (Figure 5B). Under the univariate model, the mean |t| across all 432 parcels for the 5HT-2A network (2.44) was consistently larger than for all comparator networks (5HT-1A: 1.60; 5HTT: 1.33; D1: 1.57; D2: 1.52), and the summed |t| mass was strongly biased towards 5HT-2A in all four comparisons (79-84%; donut insets). In the multivariate model the 5HT-2A mean |t| was disproportionately attenuated (1.48; a 39% reduction) compared with the comparator networks (15–30% reductions), compressing the mean |t| ratio from 1.63× to 1.25× across comparisons and reducing summed mass proportions to 57–77%. Notably, the attenuation was most severe for the comparators with the highest spatial correlation to 5HT-2A: the summed mass proportion against 5HT-1A dropped from 84% to 57%, while against 5HTT it dropped only from 79% to 77%. This asymmetry is consistent with the multivariate framework preferentially degrading receptors that share the most spatial variance with competitors, rather than uniformly attenuating all systems.

Receptor dominance analysis reinforced this pattern (Figure 5C). Under the univariate model, 5HT-2A dominated 208 of 432 parcels (48%), consistent with its established role as the primary mediator of psychedelic effects. Under the multivariate model, the largest share of parcels formerly dominated by 5HT-2A redistributed to 5HT-1A, consistent with unstable partitioning of their shared variance.

## Discussion

Molecular-enriched functional imaging methods aim to bridge neurotransmitter systems and macro-scale network dynamics. Given the inherently overlapping neurobiological organisation of neurotransmitter systems (Avery and Krichmar, 2017), spatial collinearity is inherent, yet its statistical consequences for molecular-enriched network estimation have not been systematically characterised. We demonstrate that collinearity among normative receptor maps is pervasive, scales rapidly with model complexity, and is largely unaffected by parcellation resolution. Multivariate REACT models produce less reliable molecular-enriched networks than a univariate alternative, and this degradation is driven by collinearity rather than model size per se. In an LSD pharmacoimaging dataset, the multivariate approach failed to recover the expected 5HT-2A receptor specificity. Together, these findings establish spatial collinearity as a core constraint on multivariate molecular-enriched network estimation and motivate the univariate approach as a more robust default.

The exhaustive combinatorial analysis paints a challenging picture for multivariate modelling. With four principal components accounting for 80% of the variance across 19 receptor maps, the effective dimensionality of receptor space is far lower than the number of available targets. This reflects the overlapping neurobiological organisation of neurotransmitter systems (Froudist-Walsh et al., 2023; Hansen et al., 2022), such as striatal D1/D2 co-expression (Bertran-Gonzalez et al., 2008) and hypothesised opposing roles of dopaminergic-serotonergic modulation (Boureau and Dayan, 2011). The stability of the inter-receptor correlation structure across parcellation scales is consistent with collinearity being an inherent property of macro-scale chemoarchitecture. In multivariate models, most receptor combinations exceed conventional collinearity thresholds even at the modest model sizes common in REACT analyses, and individual receptors such as D2 reach VIF values that would be considered unacceptable in any standard regression context. This suggests that previously published REACT analyses employing large receptor sets in a multivariate framework may have been operating at problematic levels of collinearity. Critically, because this collinearity reflects genuine neurobiological organisation, it cannot be resolved by restricting the receptor set: the systems most prone to co-vary are precisely those whose disentanglement most often motivates the analysis.

The key finding from the HCP analyses is not simply that reliability declines with model size, but that collinearity, rather than model complexity per se, is the dominant predictor. VIF remained highly significant after adjusting for the number of regressors, while model size added no explanatory power beyond collinearity. A negative relationship between collinearity and reliability was also evident within each level of model complexity, confirming that this is not an artefact of correlated model size and VIF. Per-receptor analyses revealed a clear neuroanatomical pattern: the most affected receptors (D2, D1, VAChT) are those with spatially concentrated distributions in the basal ganglia, where multiple neurotransmitter systems converge at high density, producing the strongest inter-map correlations and the steepest reliability declines. Conversely, receptors with more spatially distinctive cortical distributions were relatively protected. The severity of the problem is thus partially predictable from the collinearity structure, offering a prospective diagnostic for researchers selecting receptor sets.

(O’Brien, 2007), following (Belsley et al., 1980), argued that collinearity need not be treated as inherently problematic provided regression coefficients remain significant despite inflated variance, summarised in the heuristic that collinearity does not hurt so long as it does not bite. Our LSD pharmacoimaging analyses demonstrate that, even at modest levels of collinearity, it does. LSD’s subjective and neural effects are both markedly attenuated by 5HT-2A receptor antagonism with ketanserin (Becker et al., 2023; Preller et al., 2018, 2017), providing a heuristic to evaluate the two pipelines. The univariate model recovered the expected 5HT-2A dominance, with significant drug effects concentrated in the 5HT-2A network and minimal effects in comparator systems. The multivariate model, by contrast, detected no significant effects in any receptor network, despite identical data, statistical thresholds, and a relatively favourable collinearity profile. The receptor specificity analyses show that the multivariate framework disproportionately attenuated 5HT-2A effects, with dominance redistributing primarily towards 5HT-1A, the most spatially correlated comparator. This is consistent with arbitrary partitioning of shared variance between correlated predictors, eroding the signal of the pharmacologically dominant receptor.

The univariate approach is not without conceptual cost. Because each receptor is modelled independently, spatially correlated maps will yield correlated networks, and any drug effect that loads on a shared spatial pattern will appear in multiple systems. Receptor selectivity cannot therefore be established at the subject level and must emerge from the pattern of results across systems at the group level. (Kim, 2019) warns that excluding collinear variables can produce biased coefficients and that this can be potentially more serious than collinearity itself. The univariate pipeline sidesteps this objection technically but not in spirit: it does not exclude variables so much as decline to adjust for them, and each receptor’s network captures all spatially co-distributed BOLD variance rather than claiming to isolate receptor-unique contributions. More broadly, the two pipelines can be understood as providing complementary bounds on the relationship between receptor distributions and functional connectivity. The univariate approach provides an upper bound where each receptor’s network captures all spatially co-distributed BOLD variance, including variance shared with other systems. The multivariate approach (including all possible receptor/transporter maps) provides a lower bound where each receptor’s network reflects only the variance that survives partialling out all competitors. The true receptor-specific contribution lies between these extremes. However, the lower bound is only informative when it can be estimated reliably, and our results demonstrate that collinearity renders it unstable for many receptor combinations. When the lower bound is dominated by estimation noise rather than signal, the upper bound, despite its inclusiveness, provides the more interpretable estimate. In the LSD case this succeeded, with drug effects concentrated in the 5HT-2A enriched network despite substantial spatial correlation with the 5HT-1A receptor. For compounds with less well-characterised pharmacology, or where multiple systems contribute comparably, this cross-receptor reasoning will require greater care.

Although our analyses focus on REACT, the collinearity landscape is not specific to it. Any method that enters multiple PET receptor maps as simultaneous spatial regressors (Bazinet et al., 2023; Dukart et al., 2021; Hansen and Misic, 2025; Lawn et al., 2023a; Markello et al., 2022; Royer et al., 2026) faces the same underlying constraint: the effective dimensionality of receptor space is far lower than the number of available maps, and shared spatial variance among predictors will inflate uncertainty in the resulting estimates. More broadly, the standard practice of computing independent spatial correlations between brain maps and individual receptor maps is the univariate approach we advocate for here.

Several limitations should be noted. First, our analyses use normative PET maps that do not capture individual differences in receptor distributions (Hansen et al., 2026). Whether individual-level PET data would alter the collinearity landscape is an open question, though most REACT applications to date have used the same normative maps, and this reliance on group-average data is a recognised limitation across brain map contextualisation more broadly (Royer et al., 2026). Second, the exemplar LSD dataset is modest in size (N=15) and uses a single compound. Generalisability to other drugs and samples remains to be established. Third, although collinearity is stable across parcellation scales, our reliability and pharmacoimaging analyses used a single resolution, and drug-effect sensitivity may interact with parcellation scale in ways not captured here. Relatedly, parcels are not statistically independent: spatial autocorrelation in both PET and fMRI data (Alexander-Bloch et al., 2018; Leech et al., 2023; Markello and Misic, 2021; Váša and Mišić, 2022) means that increasing parcellation resolution does not proportionally increase effective sample size, limiting the extent to which finer scales can offset collinearity-induced variance inflation (O’Brien, 2007). Finally, we have not explored intermediate approaches such as regularised regression or PCA-based dimensionality reduction, because these fundamentally conflict with REACT’s analytic logic. Unlike predictive models that can be tuned to minimise out-of-sample error, REACT relies on the regression coefficients themselves as the primary scientific output. Regularisation lacks a principled optimisation target in this context and yields coefficients that no longer represent variance attributable to a specific receptor. Similarly, while PCA ensures statistical independence, it yields components that no longer correspond cleanly to neurotransmitter systems. Both remedies improve statistical stability only by sacrificing the receptor-level interpretability that motivates molecular-enriched imaging.

Altogether, we show that spatial collinearity among PET neurotransmitter maps is a pervasive property of brain chemoarchitecture that compromises multivariate molecular-enriched network estimation. We suggest that future studies employing REACT or related methods should adopt univariate pipelines as the default, report VIF diagnostics whenever multivariate models are used, and interpret multivariate results with explicit reference to the collinearity regime under which they were obtained.

## Author Contributions

**Timothy Lawn**: Conceptualization, Methodology, Software, Formal analysis, Investigation, Data curation, Visualization, Writing – original draft, Writing – review & editing. **Johan Nakuci**: Methodology, Writing – review & editing. **Steve CR Williams**: Resources, Writing – review & editing. **Federico Turkheimer**: Methodology, Writing – review & editing. **Mitul A. Mehta**: Conceptualization, Methodology, Writing – review & editing.

## Acknowledgements

Data were provided in part by the Human Connectome Project, WU-Minn Consortium (Principal Investigators: David Van Essen and Kamil Ugurbil; 1U54MH091657) funded by the 16 NIH Institutes and Centers that support the NIH Blueprint for Neuroscience Research; and by the McDonnell Center for Systems Neuroscience at Washington University. MAM receives support from the National Institute for Health and Care Research (NIHR) Maudsley Biomedical Research Centre (BRC). The views expressed are those of the author(s) and not necessarily those of the NIHR or the Department of Health and Social Care.

## Conflicts of interest statement

MAM has research funding from Nxera and Lundbeck and received in-kind contributions from Compass Pathways. He has consulted for Boehringer Ingelheim, Nxera and Curie.Bio and received speaker fees from Takeda. All other authors declare no conflicts of interest.

## Ethics Statement

This work did not collect any new data. The openly available datasets here received full ethical approval and were conducted in accordance with the revised declaration of Helsinki (2000).

## Data Availability

This work made use of publicly available datasets. The PET map data (https://github.com/netneurolab/hansen_receptors), Human Connectome Project Young Adult (HCP-YA) test-retest data (https://balsa.wustl.edu/project?project=HCP_Retest), and LSD pharmacoimaging data (https://openneuro.org/datasets/ds003059/versions/1.0.0) are all freely available for download. The Schaefer cortical atlas (https://github.com/ThomasYeoLab/CBIG/tree/master/stable_projects/brain_parcellation/Schaefer2018_LocalGlobal) and Tian subcortical atlas (https://github.com/yetianmed/subcortex) are also freely available.

## Code Availability

PET map resampling was performed using ANTs (antsApplyTransforms). Analyses were implemented in Python 3.12.4. Statistical analyses used SciPy (permutation tests, correlations, OLS regression) and statsmodels (linear mixed-effects models).

## Notes

https://openneuro.org/datasets/ds003059/versions/1.0.0

