## Supplementary materials for "Spatial collinearity constrains multivariate molecular-enriched network estimation"

---

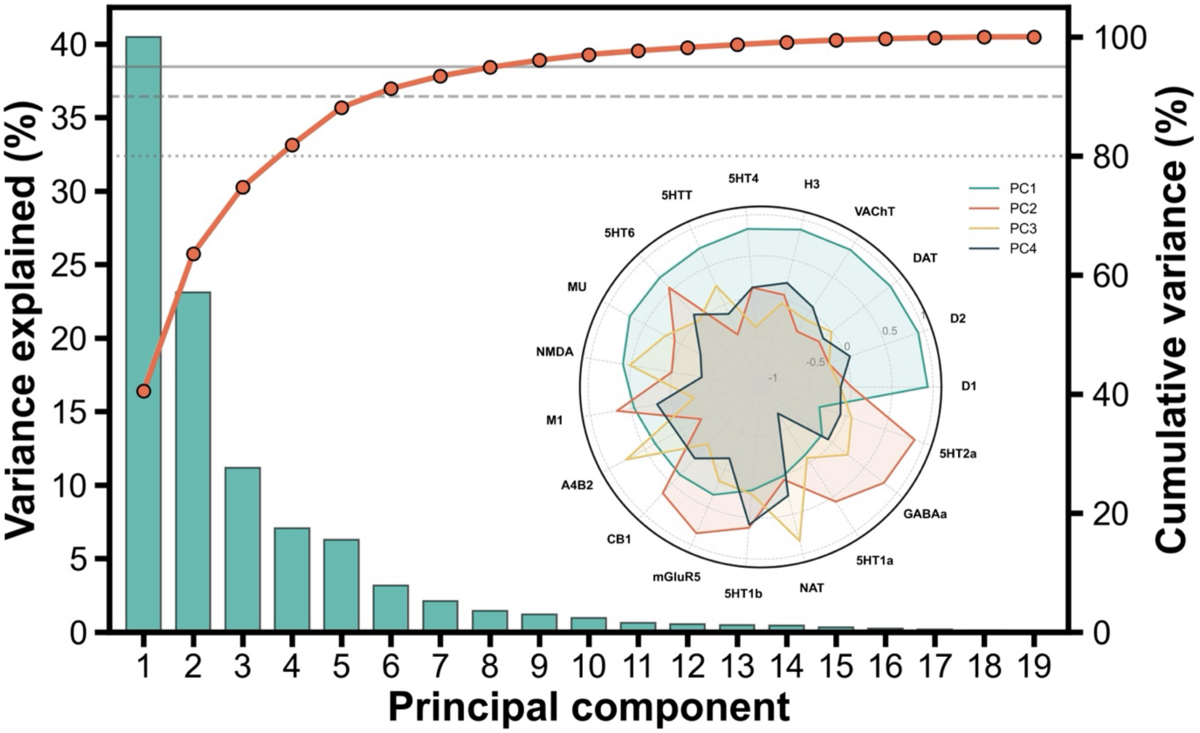


**Supplementary Figure 1 | Principal component analysis revealed the effective dimensionality of PET maps.** Bars show individual variance explained per component. Orange line shows cumulative variance. Horizontal lines indicate 80% (dotted), 90% (dashed), and 95% (solid) thresholds. Approximately four components account for 80% of the total variance across receptor and transporter maps. This highlights the spatial redundancy in the PET maps. The first four component loadings are shown for each receptor in the radar plot inlay. PC1, the dominant axis of variation, shows a subcortical-cortical contrast: maps with basal-ganglia-concentrated distributions (D1, D2, DAT, VAChT) load positively, while predominantly cortical maps (5HT-2A, GABA-A) load negatively.

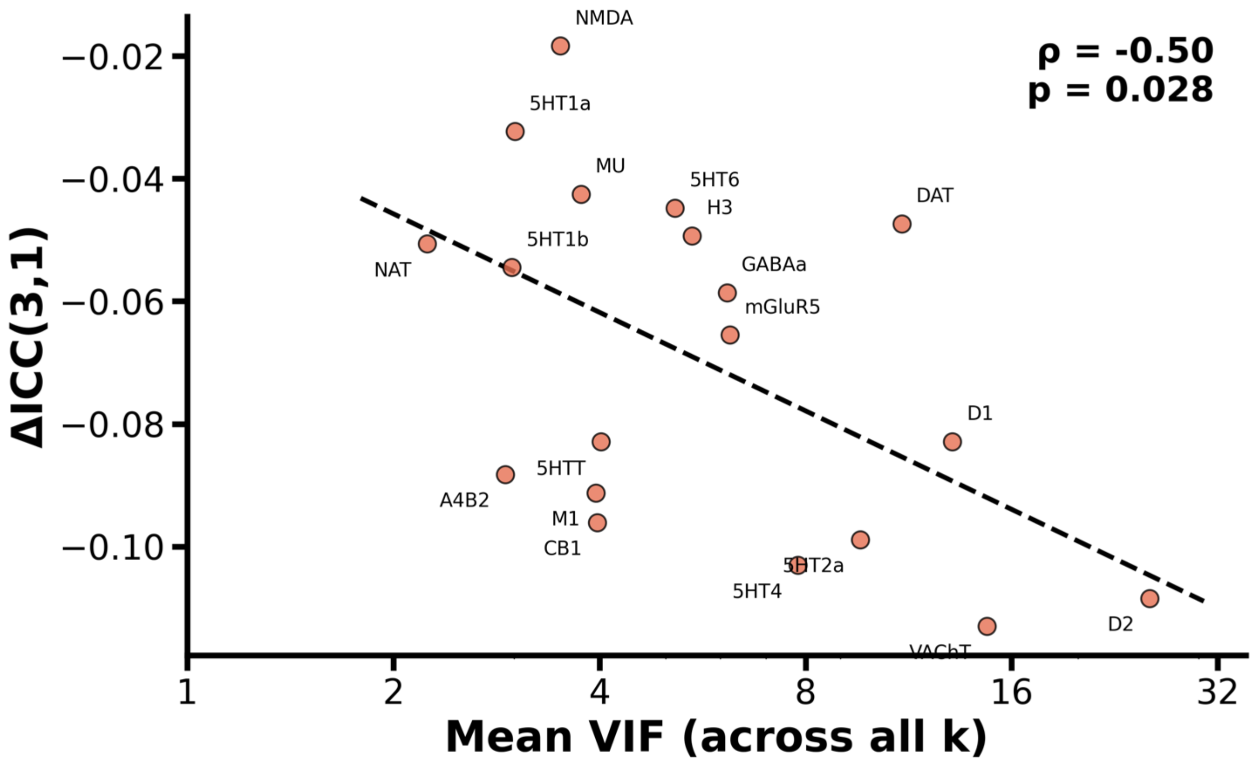


**Supplementary Figure 2 | Receptor-level correspondence between collinearity susceptibility and reliability degradation.** Each point represents one of the 19 receptors, plotting its grand mean VIF against its grand mean ΔICC, both averaged across model sizes (k=2 to 19) with each k weighted equally. These per-receptor VIF values use the same k-weighted averaging as the ΔICC, so they differ slightly from the combination-weighted ordering in Figure 2D. The dashed line shows an OLS fit on $\log_{10}$(VIF). Receptors with the most spatially overlapping distributions (e.g., D2, VAChT, D1) cluster at the high-VIF, low-ΔICC corner, while spatially distinctive receptors (e.g., NAT, 5HT-1A) are relatively protected. Spearman ρ=−0.50, p=0.028, n=19 receptors. Schaefer 400 + Tian S2 parcellation (432 parcels).
